# MEBS, a software platform to evaluate large (meta)genomic collections according to their metabolic machinery: unraveling the sulfur cycle

**DOI:** 10.1101/191288

**Authors:** Valerie De Anda, Icoquih Zapata-Peñasco, Augusto Cesar Poot-Hernandez, Luis E. Eguiarte, Bruno Contreras-Moreira, Valeria Souza

**Affiliations:** Departamento de Ecología Evolutiva, Instituto de Ecología, Universidad Nacional Autónoma de México, 70-275, Coyoacán 04510 México D.F.; Dirección de Investigación en Transformación de Hidrocarburos. Instituto Mexicano del Petróleo, Eje Central Lázaro Cárdenas, Norte 152, Col. San Bartolo Atepehuacan, 07730, México; Departamento de Ingeniería de Sistemas Computacionales y Automatización. Sección de Ingeniería de Sistemas Computacionales. Instituto de Investigaciones en Matemáticas Aplicadas y en Sistemas.; Estación Experimental de Aula Dei, Consejo Superior de Investigaciones Científicas (EEAD-CSIC), Avda. Montañana, 1005, Zaragoza 50059, Spain; Fundación ARAID, calle María de Luna 11, 50018 Zaragoza, Spain

**Keywords:** Metabolic machinery, metagenomics, omic-datasets, Pfam domains, Relative entropy, sulfur cycle, Multigenomic Entropy-based Score

## Abstract

The increasing number of metagenomic and genomic sequences has dramatically improved our understanding of microbial diversity, yet our ability to infer metabolic capabilities in such datasets remains challenging.

**FINDINGS:** We describe the Multigenomic Entropy Based Score pipeline (MEBS), a software platform designed to evaluate, compare and infer complex metabolic pathways in large ‘omic’ datasets, including entire biogeochemical cycles. MEBS is open source and available through https://github.com/eead-csic-compbio/metagenome_Pfam_score. To demonstrate its use we modeled the sulfur cycle by exhaustively curating the molecular and ecological elements involved (compounds, genes, metabolic pathways and microbial taxa). This information was reduced to a collection of 112 characteristic Pfam protein domains and a list of complete-sequenced sulfur genomes. Using the mathematical framework of relative entropy *(H’),* we quantitatively measured the enrichment of these domains among sulfur genomes. The entropy of each domain was used to both: build up a final score that indicates whether a (meta)genomic sample contains the metabolic machinery of interest and to propose marker domains in metagenomic sequences such as DsrC (PF04358). MEBS was benchmarked with a dataset of 2,107 non-redundant microbial genomes from RefSeq and 935 metagenomes from MG-RAST. Its performance, reproducibility, and robustness were evaluated using several approaches, including random sampling, linear regression models, Receiver Operator Characteristic plots and the Area Under the Curve metric (AUC). Our results support the broad applicability of this algorithm to accurately classify (AUC=0.985) hard to culture genomes (e.g., *Candidatus Desulforudis audaxviator),* previously characterized ones and metagenomic environments such as hydrothermal vents, or deep-sea sediment.

**CONCLUSIONS:** Our benchmark indicates that an entropy-based score can capture the metabolic machinery of interest and be used to efficiently classify large genomic and metagenomic datasets, including uncultivated/unexplored taxa

## Background

Over the last 15 years, the enormous advances in high-throughput sequencing technologies have revolutionized the field of microbial ecology, dramatically improving our understanding of life’s microbial diversity to an unprecedented level of detail [1-4].

Nowadays, accessing the total repertoire of genomes within complex communities by means of metagenomics is becoming a standard and routine procedure in order to attain the full insight of the diversity, ecology, evolution and functional makeup of the microbial world [5]. Furthermore, the accurate reconstruction of microbial genomes and draft-populations from environmental metagenomic studies has been shown to be a powerful approach [6-10], providing clues about the potential metabolic strategies of hard-to-culture microbial lineages by linking the functional mechanisms that support specific metabolisms with taxonomic, systematic, and ecological contexts of that lineage [8].

Despite the accelerated accumulation of large collections of metagenomic and genomic sequences, our ability to analyze, evaluate and compare complex metabolic capabilities in large-scale ‘omic’ datasets remains biologically and computationally challenging [11]. Predicting the metabolic potential is a key step in describing the relationship between a microbial community and its ecosystem function. This is largely performed by mapping the protein coding genes of ‘omic’ data onto reference pathway databases such as MetaCyc [12] or KEGG [13] based on their homology to previously characterized genes [14]. The current available methods for metabolic pathway prediction or reconstruction rely on the use of several metrics to infer the overall repertoire of metabolic pathways present in a given metagenomic dataset (e.g., MinPath [14], HUMAnN[15], PRMT [16], MetaPathways [17]).

However, due to the challenges involved in testing meaningful biological hypotheses with complex data, only a small proportion of the metabolic information derived from these datasets is eventually used to draw ecologically relevant conclusions. In this regard, most of the microbial ecology-derived ‘omic’ studies have been mainly focused on either: i) developing broad description of the metabolic pathways within a certain environment e.g., [18,19]; ii) analyzing the relative abundance of marker genes involved in several metabolic processes and in certain ecosystems (e.g., primary productivity, decomposition, biogeochemical cycling [20-24]; or iii) discovering differentially abundant, shared or unique functional units (genes, proteins or metabolic pathways) across several environmental metagenomic samples [25-27].

Therefore, in order to address some of the limitations of these methods, we propose a novel approach to reduce the complexity of targeted metabolic pathways involved in several integral ecosystem processes -- such as entire biogeochemical cycles -- into a single informative score, called Multigenomic Entropy-Based Score (MEBS). This approach is based on the mathematical rationalization of Kullback-Leibler divergence, also known as relative entropy *H’* [28]. Relative entropy has been widely applied in physics, communication theory, and statistical inference, and it is interpreted as a measure of disorder, information and uncertainty, respectively [29]. Here we use the communication theory concept of *H’* to summarize the information derived from the metabolic machinery encoded by the protein coding genes of ‘omic’ datasets. The application of this metric in biology was originally developed by Stormo and colleagues identifying binding sites that regulate gene transcription sites [30].

In order to evaluate the performance of our approach, we selected the sulfur cycle (from now on S-cycle) because this is one of the most metabolically- and ecologically complex biogeochemical cycles, but there are few studies analyzing the complete repertoire (genes, proteins, or metabolic pathways) involved in the mobilization of inorganic-organic sulfur compounds through microbial-catalyzed reactions at a planetary scale [20,31-35].

## MEBS description

MEBS (Multigenomic Entropy-Based Score, RRID: 015708) runs in Linux systems and is available at [36]. For practical purposes, the MEBS algorithm was divided into four stages summarized in Figure 1 and explained below.

**Figure 1.**
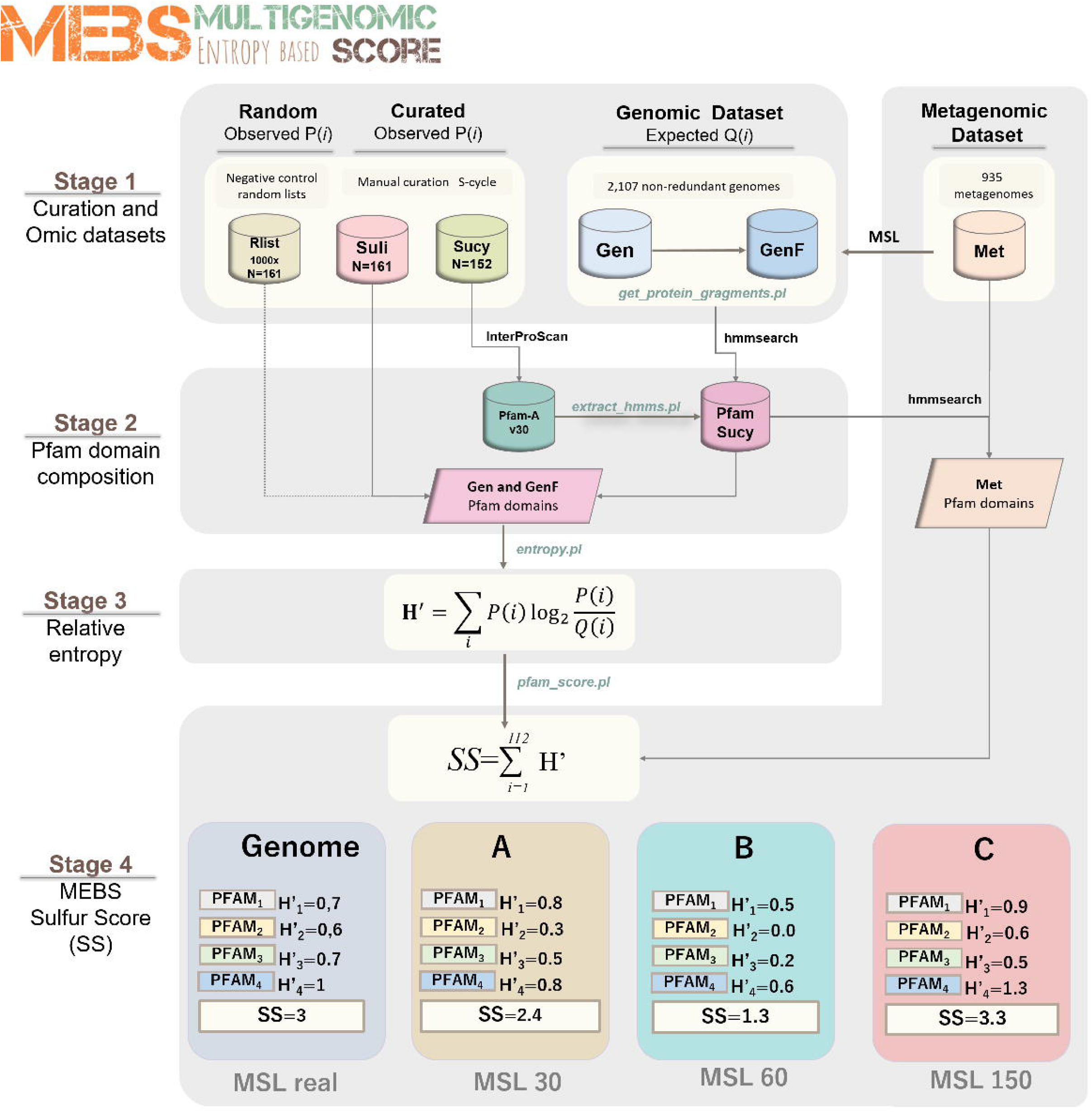
Schematic representation of the four stages of the MEBS algorithm focusing on the S-cycle. The first step consists on the systematic curation of a database containing the metabolic information of S-cycle, which is reduced to a FASTA-file of proteins involved (Sucy) and a list of 161 related microorganisms (Suli). A thousand lists of 161 random-sampled genomes were used as negative control (Rlist). The training dataset comprises 2,107 genomes (Gen), which were fragmented in different sizes by considering the Mean Size Length (MSL) of 935 metagenomes (Met). In the second stage the domain composition of Sucy proteins is obtained by scanning Pfam-A, resulting in the Pfam-Sucy database. Then, the relative entropy (/-/’) of each Sucy-Pfam domain is obtained in the third stage. Finally, the precomputed entropies in Gen and GenF are used to evaluate full-length genomic sequences (real) and metagenomic sequences of variable MSL (in this example A, B and C).

### STAGE 1: Manual curation of Sulfur cycle and ‘omic’ datasets

Sulfur taxonomic representatives. A data set comprehensively covering the currently known representatives of the S-cycle was obtained from primary literature and the MetaCyc database [12]. Each taxonomic representative (at genus or species level) was selected under the criteria of having evidence suggesting their physiological and biochemical involvement in the degradation, reduction, oxidation, or disproportionation of sulfur compounds. Then, each taxonomic representative was scanned against our Genomic dataset (see further details below), in order to obtain a list containing the completely sequenced and non-redundant genomes of the S-cycle. The resulting Sulfur list (or ‘Suli’) currently contains 161-curated genomes, and was used as the first input of the pipeline. Both the manually curated taxonomic representatives and Suli can be found in Table SI.

Random taxonomic representatives (RList). As a negative control, we generated 1000 lists of genomes that are not particularly enriched on sulfur metabolic preferences. Each list contains 161 random genomes, the same number of microorganisms included in Suli. These lists were obtained by randomly subtracting from the Genomic dataset (see below) 161 Refseq accession numbers and their corresponding names.

Metabolic pathways and genes. We gathered and classified the metabolic pathways involved in the S-cycle from the primary literature and two experimentally validated curated databases: KEGG (KEGG, RRID:SCR_012773) [13] and MetaCyc (MetaCyc, RRID:SCR_007778) [12]. All the molecular information was then combined into a single database named Sucy (for Sulfur cycle). Sucy currently contains 152 genes and 48 enzyme classification numbers annotated in the Enzyme classification [37] (Table S2). The 152 FASTA sequences of the proteins encoded by these genes were downloaded from UniProt [38] and used as the second input of the pipeline.

Genomic dataset (Gen). At the time of the analysis (December 21, 2016), a total of 4,158 genomes were available from RefSeq database [39]. For comparative genomic purposes, we removed redundancy in this large data set by using the Web interface [40] described in [41]. As phylogenomic distance measure, we used a modified version of the Genomic Similarity Score defined as GSSb in [41]; we selected the most tolerant threshold of 0.95 (so as not to drop many sequenced genomes) and default parameters, resulting in 2,107 clusters containing similar genomes, ordered by size (largest to smallest). Then, the largest genome representative for each group was searched in the NCBI genome assembly summary file [42] and downloaded from the NCBI FTP site [43].

Metagenomic dataset (Met). We used the Meta Genome Rapid Annotation using Sub-system Technology server (MG-RAST, RRID:SCR_004814) [44] to download metagenomes that: i) were publicly available; ii) contained associated metadata; and iii) had been isolated from well-defined environments (i.e., rivers, soil, biofilms), discarding host associated microbiome sequences (i.e., human, cow, chicken). In addition we also included 35 unpublished metagenomes derived from sediment, water and microbial mats from Cuatro Ciénegas, Coahuila (CCC), Mexico. The latter were also submitted and annotated in the MG-RAST server, and will be described in depth elsewhere. The resulting collection of 935 FASTA files (≈500 GB), containing gene-called protein sequences (MG-RAST stage 350), were downloaded from the RESTful MG-RAST API (http://api.metagenomics.anl.gov/api.html). While these metagenomes were evaluated and scored in STAGE 4, they were also analyzed to estimate their mean sequence length, considering that the fragmented nature of metagenomic sequences would have an impact on homology detection, depending on the length of the reads [45,46]. Therefore, we measured the Mean Size Length (MSL) of the peptide sequences of the 935 metagenomes in Met and the 152-curated proteins in Sucy, which are summarized in Figure S1. It was observed that the MSL of Met varies broadly, with a majority of metagenomic peptides with MSL ≤ 30 aa, and that Sucy proteins range from 49 to 1,020 aa, with MSL=349 aa. According to this distribution, the metagenomes in Met were grouped into seven well-defined categories: MSL≤30, ≤60, ≤100, ≤150, ≤200, ≤250, ≤300 aa.

Fragmented genomic dataset (GenF). In order to simulate the observed variability of MSL across metagenomes, protein sequences encoded in the genomic dataset (Gen, containing 2,107 genomes) were *in silico* sheared with Perl script *get_protein_fragments.pl* into the seven MSL categories defined above (30 to 300). This produced the GenF dataset, which currently requires up to 104GB of disk space.

### STAGE 2: Domain composition of the input proteins

The annotation of protein domains in Sucy was conducted using Interproscan 5.21-60.0 [47] against databases Pfam-A v30 (Pfam, RRID:SCR_004726) [48], TIGRFAM v13 (JCVI TIGRFAMS, RRI D:SCR_005493) [49] and Superfamily v1.75 (SUPERFAMILY, RRID:SCR_007952) [50]. Then, the Hidden Markov Models (HMMs) from matched Pfam domains (n=112) were extracted from Pfam-A using script *extract_hmms.pl.* These selected HMMs were subsequently scanned against the Genomic, Genomic Fragmented and Metagenomic datasets (from now on ‘omic’ datasets, see subsequent stages) using HMMER 3.0 *hmmsearch*--cut_ga option [51].

### STAGE 3: Relative entropy and its use in detecting informative domains

In order to detect protein domains enriched among sulfur-based microorganisms (Suli), we used a derivative of the Kullback-Leibler divergence [28] — also known as relative entropy *H’(i) —* to measure the difference between probabilities *P* and Q (see Eq. 1 below). In this context, *P(i)* represents the frequency of protein domain / in the 161 Suli genomes (observed frequency), while *Q(i)* represents its frequency in the 2,107 genomes in Gen (expected frequency). The script to compute the entropy *(entropy.pl)* requires the list of the genomes of interest (Suli) and the tabular output file obtained in from the scanning of Gen and GenF against Pfam-Sucy database. The obtained values of *H’* (in bits) capture to what extent a given Pfam domain informs about the metabolism of interest. In this case, domains with *H’* values close or greater than one, correspond to the most informative Pfam domains (enriched among S-based genomes), whereas low *H’* values (close to zero) indicate non-informative ones. Negative values correspond to those observed less than expected.

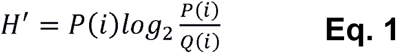

As a negative control, the *H’* of the 112 Pfam domains were recalculated in both Gen and GenF datasets, but replacing Suli with 1,000 equally sized lists of random-sampled genomes (Rlist). We evaluated the impact of the MSL in the computed entropy values using Gen and GenF. First, we focused on detecting informative Pfam domains that could be used as possible molecular marker genes in variable length, metagenomic sequences. Specifically, we looked for domains displaying stable *H’* values across both Gen and GenF by using the script *plot_cluster_comparison.py,* which implements the following methods: K-Means, Affinity propagation, Mean-shift Spectral, Ward hierarchical, Agglomerative, DBSCAN and Birch. All of these are part of the scikit-learn Machine Learning Python module [52].

### STAGE 4: Final score, interpretation, properties and benchmark

Peptide sequences from a given genome or metagenome of interest are evaluated by first scanning their Pfam domains and then producing a final score, defined as the sum of the precomputed entropies of matched S-related Pfam domains (see Equation 2). This score (Sulfur Score ‘SS’ in our case) summarizes the information content of the metabolic machinery of interest. In this context, informative sulfur protein domains would contribute to higher SS, whereas non-informative ones would decrease it. This is an extension of procedures originally developed for the alignment of DNA and protein motifs, in which individual positions are independent and additive, and can be simply summed up to obtain the total weight or information content [30]. Instead of aligning sequences, in our context we added up the entropy values of the Pfam domains matched in a given ‘omic’ sample (resulting from scanning the sample of interest against Pfam-Sucy), from which a total weight (SS) is computed by using script *pfam_score.pl.*

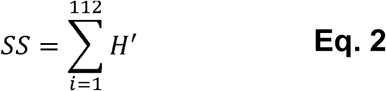

Datasets in which the majority of informative S-cycle protein domains are represented will yield a high SS; in contrast, low SS values should be expected if proteins involved in the S-cycle are not particularly enriched.

MSL. As the calculation of the SS depends on the MSL of the omic sample of interest, script *pfam_score.pl* supports option-size, in amino acid residues (aa). In this way, appropriate precomputed *H’* values for Pfam domains can be selected to produce the final score. Currently 30, 60, 100, 150, 200, 250, 300 and real sizes are supported.

Metabolic pathway completeness and KEGG visualization. The presence-absence patterns of Pfam domains belonging to particular pathways can be exploited to compute metabolic completeness. This optional task is invoked with parameter-keggmap and a TAB-separated file mapping Pfam identifiers to KEGG Orthology entries (KO numbers) and the corresponding pathway in Sucy (see Table S3). To compute completeness, the total number of domains involved in a given pathway (i.e., sulfate reduction, sulfide oxidation) must be retrieved from the Sucy database (See Table S2). Then, the protein domains currently present in any given sample are divided by the total number of domains in the pre-defined pathway. The script produces: i) a detailed report of the metabolic pathways of interest; and ii) a list of KO numbers with Hex color codes, corresponding to KO matches in the omic sample, which can be exported to the KEGG Mapper - Search & Color Pathway tool [53] (see Figure S2).

Properties and performance of SS. Since the outcome of the final score (SS) largely depends on the list of microorganisms involved in the metabolism of interest (in our case Suli) and the Pfam domains found in the input protein sequences (n=112), we evaluated its robustness and reproducibility with several approaches. First, we compared our results with a benchmark performed three years ago in which we used Pfam-A v27 (instead of version 30), a genomic dataset containing 1,528 non-redundant genomes (579 less genomes than our current Genomic dataset), and an input list of 156 genomes of interest (five less that our current Suli). Second, SS estimates were compared with scores obtained by randomly selecting ≈50% of the 112 Pfam domains with both Gen and Met. This analysis was performed a thousand times with *pfam_score.pl*-random. Third, we benchmarked the predictive capacity of the SS in order to accurately classify genomes of S-related organisms (Suli, n=161, positive instances), in contrast with a larger set of non-redundant genomes (Gen - Suli, n=1.946, negative instances). Therefore, we computed the True Positive Rates (TPR), False Positive Rates (FPR), Receiver Operating Characteristic (ROC) plots and the resulting Area Under the Curve (AUC) using the scikit-learn module described in [52].

## Results and discussion

We present MEBS a new open source software to evaluate, quantify, compare, and predict the metabolic machinery of interest in large ‘omic’ datasets. The pipeline includes four stages. The first one consists on the systematic and targeted acquisition of the molecular and ecological information describing the metabolism of interest, represented by a list of curated microorganisms and a FASTA file of proteins involved in that metabolic network. In the second stage, the domain composition of the curated proteins is evaluated. Then, the domains enriched among the microorganisms of interest are identified by using the mathematical framework of the relative entropy (/-/’, third stage). Finally, the summation of the entropy of individual Pfam domains in a given genome or metagenomic dataset yields the final score (see Figure 1).

To test the applicability of this approach, we evaluated the metabolic machinery of the S-cycle. Due to its multiple redox states and its consequences on microbiological and geochemical transformations, S-metabolism can be observed as a complex metabolic machinery, involving a myriad of genes, enzymes, organic substrates and electron carriers, which largely depend on the surrounding geochemical and ecological conditions. For these reasons, the complete repertory involved in the metabolic machinery of S-cycle has remained underexplored despite the massive data produced in ‘omic’ experiments. Here, we performed an integral curation effort to describe all the elements involved in the S-cycle and then used, as explained in the following sections, to score genomic and metagenomic datasets in terms of their Sulfur relevance.

### Manual curation: the complex metabolic machinery of the Sulfur cycle

In order to integrate the complete biogeochemical S-cycle, we manually curated and modeled the major processes involved in the mobilization and use of S-compounds through Earth biosphere. This effort resulted in two comprehensive databases. The first one includes most of the known microorganisms (with and without complete genomes) described in the literature to be closely involved in the S-cycle (Table S1). In this database, we included representative taxa from the following metabolic sulfur guilds: i) chemolithotrophic, colorless sulfur bacteria (CLSB: 24 genera); ii) anaerobic phototrophs, purple sulfur bacteria (PSB:25 genera), and green sulfur bacteria (GSB:9 genera); iii) sulfate reducing bacteria (SRB: 40 genera); and iv) deep-branch sulfur hyperthermophilic microorganisms, such as elemental sulfur reducing (SRM:19 genera) and oxidizers (SO:4 genera). From all the microorganisms described to be involved in the S-cycle, at the time of the analysis, a total of 161 were found to be completely sequenced and non-redundant genomes, and were used as the first input of the pipeline (Suli).

The second database (Sucy) contains genes, proteins, and pathways with experimental evidence linking them to the S-cycle. To compile this database, we first gathered the most important S-compounds derived from biogeochemical processes and biological catalyzed reactions. Then we classified each S-compound according to their chemical and thermodynamic nature (Gibbs free energy of formation, GFEF). Finally, we classified weather each compound can be used as a source of carbon, nitrogen, energy or electron donor, fermentative substrate, or terminal electron acceptor in respiratory microbial processes. The schematic representation of the manual curated effort summarizing the complexity of the sulfur biogeochemical cycle in a global scale is shown in Figure 2.

**Figure 2.**
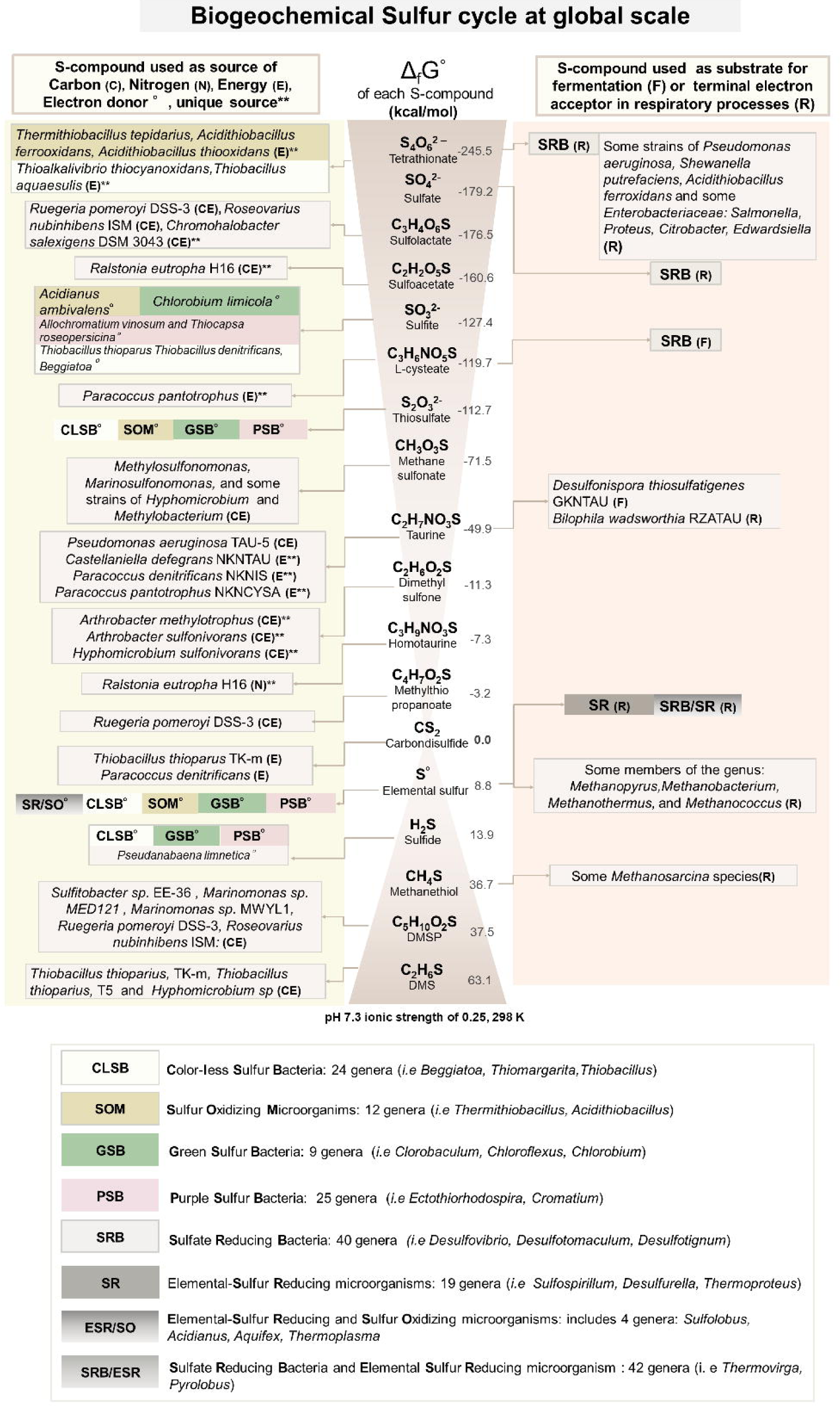
Sulfur cycle at global scale. The most important organic and inorganic S-compounds derived from biogeochemical processes are arranged according to the Standard Gibbs free energy of formation described in Caspi et al., (2012). The left column indicates whether specific microorganisms are able to use those S-compounds, as a source of Carbon (C), Nitrogen (N), Energy (E) or Electron donors (°). Double asterisks indicate whether the S-compound is used as sole source, of C, N, or E. The corresponding electron acceptors in redox-coupled reactions using the S-compound as electron donor are not shown. The right column indicates whether the S-compound is used as fermentative substrate (F) or terminal electron acceptor in respiratory processes (R). Colored boxes summarize the metabolic guilds involved in the metabolism of S-compounds, in oxidation (i.e., CLSB, SOM, PSB, and GSB) or reduction (SR, SRB) processes. The complete list of S-based microorganisms (Suli) is found in Table S1. Figure based on annotations from MetaCyc [12],

Once we selected the microorganisms, genes, and biogeochemical processes involved, we systematically divided the metabolic machinery of the S-cycle into 28 major metabolic pathways described in Table 1. In general terms we included pathways involved in: i) the oxidation/reduction of inorganic S-compounds, used as source of energy, electron donor or acceptor (P1-P7, P11 and P20 and P21); ii) the degradation of organic S-compounds, such as aliphatic sulfonates, sulfur amino acids, and organosulfonates (P8-P10, P12-P19, P22,P23,P27); iii) the methanogenesis from methylated thiols, such as dimethyl sulfide DMS (P24), metylthio-propanoate (P25) and methanethiol(P26), which are generated in nature by different biogeochemical processes [12]; and finally, iv) the biosynthesis of sulfolipids (SQDG) (P28), because it has been observed that some bacteria living in S-rich and P-lacking environments are able to synthetize sulfolipids, instead of phospholipids, in the membrane as an adaptation to the selective pressures of these particular environments [54]. The synthetic pathway P29 is explained in further detail in the next sections (Table 1).

**Table 1.** Metabolic pathways of global biogeochemical S-cycle

| Pathway number | Metabolism<sup>a</sup> | Chemical process<sup>b</sup> | Sulfur compound | Type<sup>c</sup> | Chemical formula | Source<sup>d</sup> | Number of Pfam domains<sup>e</sup> |
| --- | --- | --- | --- | --- | --- | --- | --- |
| P1 | DS | O | Sulfite | I | SO<sub>3</sub><sup>2-</sup> | E | 9 |
| P2 | DS | O | Thiosulfate | I | S<sub>2</sub>O<sub>3</sub><sup>2-</sup> | E | 10 |
| P3 | DS | O | Tetrathionate | I | S<sub>4</sub>O<sub>6</sub><sup>2-</sup> | E | 2 |
| P4 | DS | R | Tetrathionate | I | S<sub>4</sub>O<sub>6</sub><sup>2-</sup> | E | 17 |
| P5 | DS | R | Sulfate | I | SO<sub>4</sub><sup>2-</sup> | E | 20 |
| P6 | DS | R | Elemental sulfur | I | S<sup>0</sup> | E | 20 |
| P7 | DS | D | Thiosulfate | I | S<sub>2</sub>O<sub>3</sub><sup>2-</sup> | E | 9 |
| P8 | DS | O | Carbon disulfide | O | CS<sub>2</sub> | E | 1 |
| P9 | A | DE | Alkanesulfonate | O | CH<sub>3</sub>O<sub>3</sub>SR | S | 5 |
| P10 | A | R | Sulfate | I | SO<sub>4</sub><sup>2-</sup> | S | 20 |
| P11 | DS | O | Sulfide | I | H<sub>2</sub>S | E/S | 29 |
| P12 | A | DE | L-cysteate | O | C<sub>3</sub>H<sub>6</sub>NO<sub>5</sub>S | C/E | 1 |
| P13 | A | DE | Dimethyl sulfone | O | C<sub>2</sub>H<sub>6</sub>O<sub>2</sub>S | C/E | 3 |
| P14 | A | DE | Sulfoacetate | O | C<sub>2</sub>H<sub>2</sub>O<sub>5</sub>S | C/E | 2 |
| P15 | A | DE | Sulfolactate | O | C<sub>3</sub>H<sub>4</sub>O<sub>6</sub>S | C/S | 14 |
| P16 | A | DE | Dimethyl sulfide | O | C<sub>2</sub>H<sub>6</sub>S | C/S | 16 |
| P17 | A | DE | Dimethylsulfoniopropionate | O | C<sub>5</sub>H<sub>10</sub>O<sub>2</sub>S | C/S/E | 12 |
| P18 | A | DE | Methylthiopropionate | O | C<sub>4</sub>H<sub>7</sub>O<sub>2</sub>S | C/S | 7 |
| P19 | A | DE | Sulfoacetaldehyde | O | C<sub>2</sub>H<sub>3</sub>O<sub>4</sub>S | C/S | 7 |
| P20 | DS | O | Elemental sulfur | I | S<sup>0</sup> | C/S/E | 7 |
| P21 | DS | D | Elemental sulfur | I | S<sup>0</sup> | C/S/E | 1 |
| P22 | A | DE | Methanesulfonate | O | CH<sub>3</sub>O<sub>3</sub>S | C/S/E | 7 |
| P23 | A | DE | Taurine | O | C<sub>2</sub>H<sub>7</sub>NO<sub>3</sub>S | C/S/E | 11 |
| P24 | DS | M | Dimethyl sulfide | O | C<sub>2</sub>H<sub>6</sub>S | C | 1 |
| P25 | DS | M | Methylthio-propionate | O | C<sub>4</sub>H<sub>7</sub>O<sub>2</sub>S | C | 1 |
| P26 | DS | M | Methanethiol | O | CH<sub>4</sub>S | C | 1 |
| P27 | A | DE | Homotaurine | O | C<sub>3</sub>H<sub>9</sub>NO<sub>3</sub>S | N | 1 |
| P28 | A | B | Sulfolipid | O | SQDG |  | 4 |
| P29 |  |  | Markers |  | Markers |  | 12 |

<sup>a</sup> Metabolism: Assimilative (A) inorganic compounds are reduced during biosynthesis; Dissimilative (DS) inorganic compounds used as electron acceptors in energy metabolism. A large amount of electron acceptor is reduced and the reduced product is secreted.
<sup>b</sup> Chemical Process: Oxidation (O): Reduction (R), Degradation (DE), Biosynthesis (B), Methanogenesis (M), Disproportionation (D).
<sup>c</sup> Compound Type: Organic (O): sulfur atoms with covalent bonds to carbon atoms. Inorganic (I): sulfur compounds with non-carbon atoms.
<sup>d</sup> Source: sulfur compound used as source of energy (E), sulfur (S), carbon (C), nitrogen (N).
<sup>e</sup> Number of Pfam domains belonging to each metabolic pathway described in Sucy (Table S2)

After the comprehensive metabolic inventory was compiled, we linked all the elements in a single network representation of the S-metabolic machinery (Figure 3). To the best of our knowledge, this is the first molecular reconstruction of the cycle that considers all the sulfur compounds, genes, proteins and the corresponding enzymatic steps resulting into higher order molecular pathways. The latter representation also highlights the interconnection of pathways in terms of energy flow and the interplay of the redox gradient (organic/inorganic) of the intermediate compounds that act as key axes of organic and inorganic reactions (e.g., sulfite).

**Figure 3.**
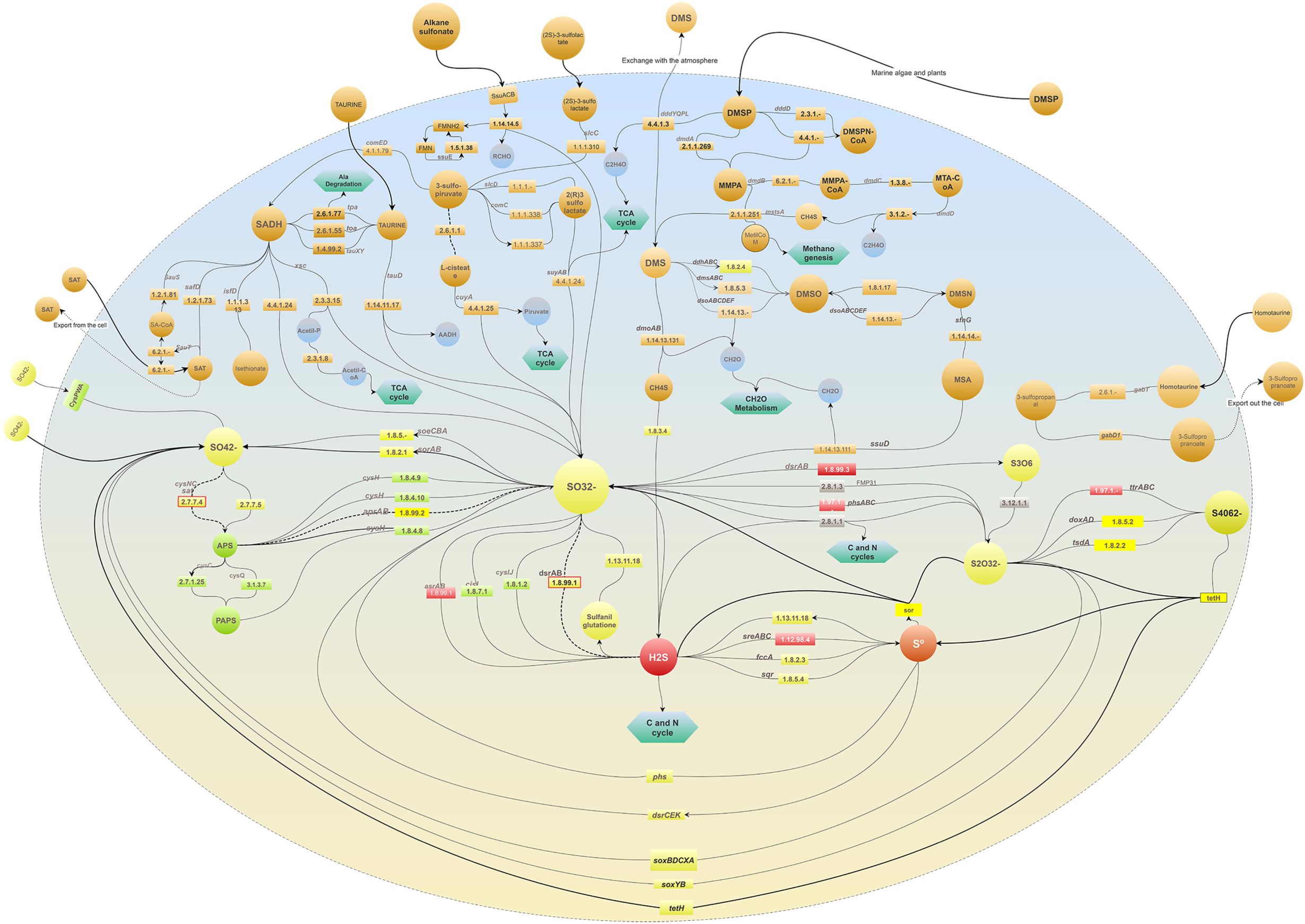
Comprehensive network representation of the machinery of the biogeochemical S-cycle in a single cell. The 28 molecular pathways involved in the metabolism of sulfur compounds described in Table 1 are included. The enzymatic steps are depicted as rectangles followed by arrows indicating the direction of the reaction. Green hexagons represent metabolic links to other metabolisms. Bold dashed arrows indicate bidirectional reactions. Inorganic S-compounds have been arranged according to their reduction potential, from the most oxidized (yellow) to the most reduced (red) compounds. Grey rectangles indicate enzymes acting in disproportionation processes in which a reactant is both oxidized and reduced in the same chemical reaction, forming two separate compounds. Input biogeochemical S-compounds are shown outside and connected with bold arrows. Dashed arrows indicate S-compounds excreted out of the cell. The upper half of the modeled cell depicts the processes involved in the use of organic S-compounds (orange circles) found in natural environments and used as source of carbon, sulfur and/or energy in several aerobic/anaerobic strains described in Figure 2.

### Annotation of Pfam domains within Sulfur proteins

Our approach requires the detection of structural and evolutionary units, also known as domains, in the curated list of protein sequences involved in the metabolism of interest (S-cycle in this case). The annotation of protein domains against the Pfam-A database resulted in a total of 112 domains identified in 147 proteins (out of 152). These 112 domains constitute the Pfam-Sucy database and represent all the pathways listed in Table 1. Two other protein family databases were tested (TGRFAM and Superfamily), but the number of proteins with positive matches was lower than with Pfam (57 and 137, respectively) and thus were not further considered.

### Preparation of omic datasets: Gen, GenF and Met

The genomic dataset required for computing domain entropies (Gen) was obtained from public databases, as explained above in MEBS Description. A fragmented version of Gen, called GenF, was generated by considering the Mean Size Length (MSL) distribution of metagenomic sequences (Figure S1).

In order to benchmark MEBS with real environmental metagenomic samples, a collection of 900 public metagenomes was obtained from MG-RAST, to which we added 35 metagenomes sampled from an ultra-oligotrophic shallow lake in México (CCC). Altogether, these 935 metagenomes set up the Met dataset.

### Using the relative entropy to recognize S-cycle domains and candidate markers

The next stage consists on the quantitative detection of informative domains (enriched among organism in Suli), by computing its relative entropy *(H’)* using Equation 1. The occurrences of each of the 112 Pfam domains in Suli and the genomic datasets were taken as observed and expected frequencies, respectively. Figure 4A summarizes the computed *H’* values in real (Gen) and fragmented genomic sequences of increasing size (GenF). The results indicate that only a few Pfam domains are equally informative regardless of the length of sequences. When *H’* values inferred from real, full-length proteins are compared to those of fragmented sequences, it can be seen that shorter sequences (MSL 30 & 60 aa) yield larger entropy differences than sequences of length > 100 aa (see in Figure 4B). Therefore, in order to shortlist candidate marker genes we selected those Pfam domains displaying constant, high mean *H’* values in Gen and GenF, low *H’* standard deviation (std) and a clear separation from the random distribution.

**Figure 4.**
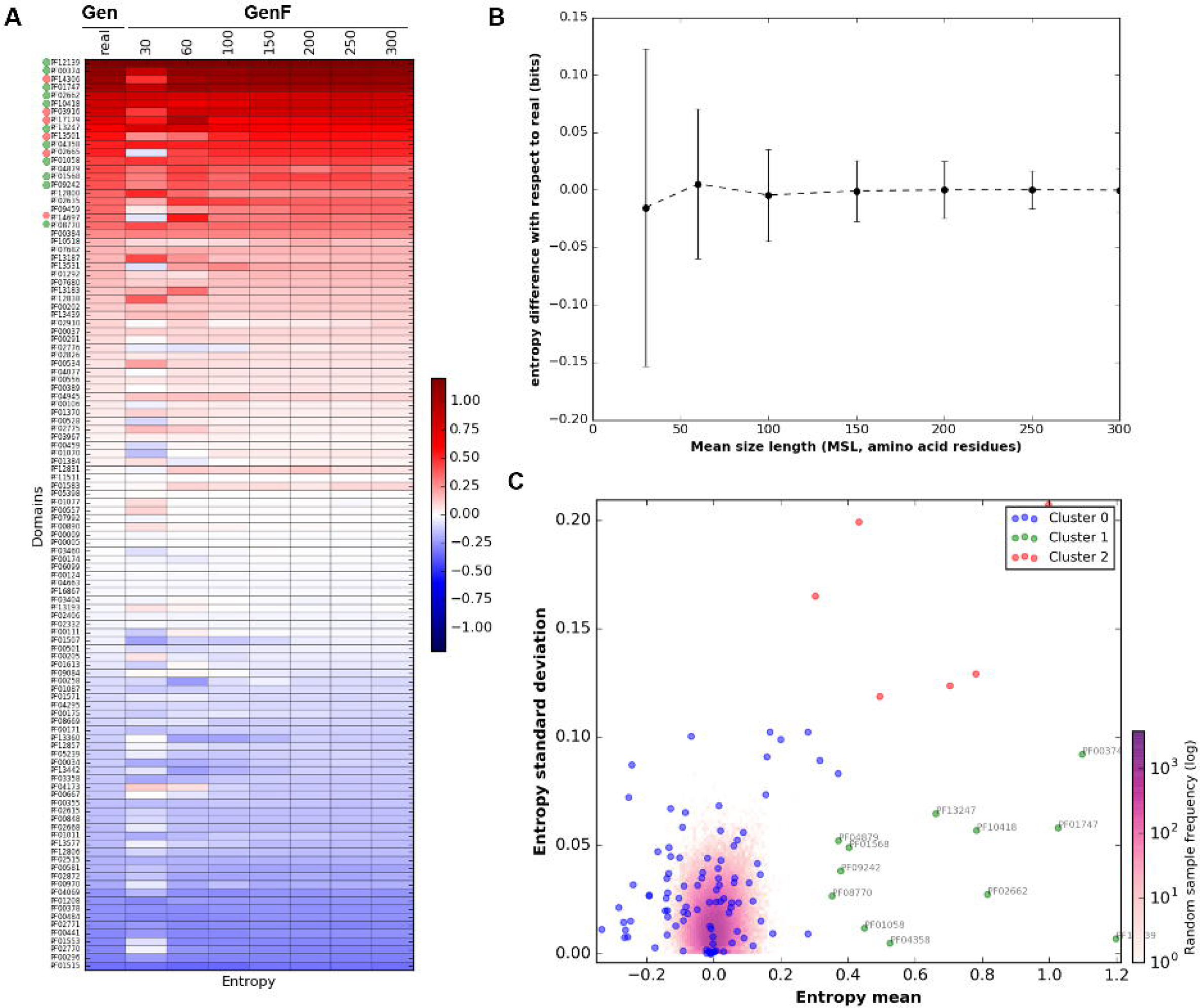
Entropy values of Sulfur-derived protein domains. A) Heatmap showing the entropy values (*H’)* of the 112 Pfam domains identified in proteins curated in SuCy. B) Difference between entropies estimated from sizes categories of growing peptide size (GenF) and the real values measured within complete genomes (Gen). Error bars show standard deviations. Both graphs were obtained with script *p!ot_entropy.py.* Clustering of the Pfam relative entropies obtained in Gen and GenF produced with the Ward method. Log frequency of the entropy values computed in the random test is colored in purple (see scale bar). Cluster 0 (blue) groups protein domains with low relative entropy that overlap with the random distribution. Cluster 1 (green) includes the Pfam domains that fulfill the requirements to be used as molecular markers (high H’ and low standard deviation, std). Red dots (cluster 2) correspond to Pfam domains with high *H’* and std. The cluster was produced with script *F_meanVSstd.py*

We tested several clustering methods, summarized in Figure S3, with Ward and Birch performing best in grouping together informative protein domains with low std. However, the Ward classification was eventually selected as Birch failed to include a few Pfam domains relevant in the S-cycle (see Figure S4). By using Ward method, three well-defined clusters of Pfam domains were generated, as observed in Figure 4C. Cluster 0 included 94 domains containing H’ values ranging from [-0.4, 0.4] and overlapping with the values obtained in the negative control explained in the next section. Cluster 1 consistently grouped together 12 Pfam domains listed in Table 2 with high entropy and low std, and can therefore be proposed as molecular markers in metagenomic sequences of variable length. Among the proposed marker domains are APS-Reductase (PF12139: H’=1.2), ATP-sulfurilase (PF01747: H’=1.03) and DsrC (PF04358: H’=0.52), key protein families in metabolic pathways involved in both sulfur oxidation/reduction processes. Finally, cluster 2 includes Pfam domains displaying high entropy values and high std, such as the PUA-like domain (PF14306: H’=1). We presume that domains within this cluster are also key players in S-metabolism; however, their high std makes them unsuitable for markers, particularly with metagenomic sequences of variable MSL. We suggest that further analyses will be required to test the implication in S-energy conservation processes of proteins containing domains such as PF03916, PF02665 or PF14697 (see complete list in Table S4).

**Table 2.** Informative Pfam domains with high *H’* and low std. Novel proposed molecular marker domains in metagenomic data of variable MSL

| Pfam ID<br>( Suli<br>ocurrences) | $H'$<br>mean | $H'$<br>std | Description |
| --- | --- | --- | --- |
| PF12139<br>58/161 | 1.2 | 0.01 | **Adenosine-5'-phosphosulfate reductase beta subunit:** Key protein domain for both sulfur oxidation/reduction metabolic pathways. Has been widely studied in the dissimilatory sulfate reduction metabolism. In all recognized sulfate-reducing prokaryotes, the dissimilatory process is mediated by three key enzymes: Sat, Apr and Dsr. Homologous proteins are also present in the anoxygenic photolithotrophic and chemolithotrophic sulfur-oxidizing bacteria (CLSB, PSB, GSB), in different cluster organization [35]. |
| PF00374<br>135/161 | 1.1 | 0.09 | **Nickel-dependent hydrogenase:** Hydrogenases with S-cluster and selenium containing Cys-x-x-Cys motifs involved in the binding of nickel. Among the homologues of this hydrogenase domain, is the alpha subunit of the sulfhydrogenase I complex of *Pyrococcus furiosus*, that catalyzes the reduction of polysulfide to hydrogen sulfide with NADPH as the electron donor [55]. |
| PF01747<br>103/161 | 1.03 | 0.06 | **ATP-sulfurylase:** Key protein domain for both sulfur oxidation and reduction processes. The enzyme catalyzes the transfer of the adenyl group from ATP to inorganic sulfate, producing adenosine 5'-phosphosulfate (APS) and pyrophosphate, or the reverse reaction [56]. |
| PF02662<br>62/161 | 0.82 | 0.03 | **Methyl-viologen-reducing hydrogenase, delta subunit:** Is one of the enzymes involved in methanogenesis and encoded in the mth-flp-mvh-mrt cluster of methane genes in *Methanothermobacter thermautotrophicus*. No specific functions have been assigned to the delta subunit [48]. |
| PF10418<br>122/161 | 0.78 | 0.06 | **Iron-sulfur cluster binding domain of dihydroorotate dehydrogenase B:** Among the homologous genes in this family are *asrA* and *asrB* from *Salmonella enterica enterica serovar Typhimurium*, which encode 1) a dissimilatory sulfite reductase, 2) a gamma subunit of the sulfhydrogenase I complex of *Pyrococcus furiosus* and, 3) a gamma subunit of the sulfhydrogenase II complex of the same organism [12]. |
| PF13247<br>149/161 | 0.66 | 0.06 | **4Fe-4S dicluster domain:** Homologues of this family include: 1) DsrO, a ferredoxin-like protein, related to the electron transfer subunits of respiratory enzymes, 2) dimethylsulfide dehydrogenase $\beta$ subunit (ddhB), involved in dimethyl sulfide degradation in *Rhodovulum sulfidophilum* and 3) sulfur reductase FeS subunit (sreB) of *Acidianus ambivalens*, involved in the sulfur reduction using $H_2$ or organic substrates as electron donors [12]. |
| PF04358<br>73/161 | 0.52 | 0 | **DsrC like protein:** DsrC is present in all organisms encoding a dsrAB sulfite reductase (sulfate/sulfite reducers or sulfur oxidizers). The physiological studies suggest that sulfate reduction rates are determined by cellular levels of this protein. The dissimilatory sulfate reduction couples the four-electron reduction of the DsrC trisulfide to energy conservation [57]. DsrC was initially described as a subunit of DsrAB, forming a tight complex; however, it is not a subunit, but rather a protein with which DsrAB interacts. DsrC is involved in sulfur-transfer reactions; there is a disulfide bond between the two DsrC cysteines as a redox-active center in the sulfite reduction pathway. Moreover, DsrC is among the most highly expressed sulfur energy metabolism genes in isolated organisms and meta- transcriptomes (Santos et al., 2015). |
| PF01058<br>158/161 | 0.45 | 0.01 | **NADH ubiquinone oxidoreductase, 20 Kd subunit:** Homologous genes are found in the delta subunits of both sulfhydrogenase complexes of *Pyrococcus furiosus* [12]. |
| PF01568<br>156/161 | 0.4 | 0.05 | **Molydopterin dinucleotide binding domain:** This domain corresponds to the C-terminal domain IV in dimethyl sulfoxide (DMSO) reductase [48]. |
| PF09242<br>39/161 | 0.38 | 0.04 | **Flavocytochrome c sulphide dehydrogenase, flavin-binding:** Enzymes found in S-oxidizing bacteria such as the purple phototrophic bacteria *Chromatium vinosum* [48]. |
| PF04879<br>151/161 | 0.37 | 0.05 | **Molybdopterin oxidoreductase Fe4S4 domain:** Is found in a number of reductase/dehydrogenase families, which include the periplasmic nitrate reductase precursor and the formate dehydrogenase alpha chain, i.e., *Wolinella succinogenes* polysulfide reductase chain. *Salmonella typhimurium* thiosulfate reductase (gene phsA). |
| PF08770<br>45/161 | 0.35 | 0.03 | **Sulphur oxidation protein SoxZ:** SoxZ sulfur compound chelating protein, part of the complex known as the Sox enzyme system (for sulfur oxidation) that is able to oxidize thiosulfate to sulfate with no intermediates in *Paracoccus parantropus* [12]. |

### Is the entropy affected by the input list of microorganisms? Negative control test

In order to evaluate to what extent the *H’* values depend on the curated list of microorganisms, we performed a negative control by replacing Suli in 1,000 lists of randomly-sampled genomes and used them to compute the observed frequencies (see Equation 1). As expected, there was a clear difference between both *H’* estimates (see Figure S5). In particular, entropy values derived from the random test were found to be approximately symmetric and consistently low among the GenF size categories (compared with the real values), yielding values of -0.09, and 0.1 as 5% and 95% percentiles, respectively (Table S5).

### Sulfur Score and its predictive capacity to detect S-microbial players in a large genomic dataset

To test whether Pfam entropies can be combined to capture the S-metabolic machinery in ‘omic’-samples, we calculated the final MEBS score, called in this case Sulfur Score (SS). We computed the SS on each of the 2,107 non-redundant genomes in Gen with script *score_genomes.sh.* The individual genomes along with their corresponding SS values and taxonomy according to NCBI are found in Table S6.

For evaluation purposes, we classified and manually annotated all the genomes in Gen according to their metabolic capabilities. First, we identified the 161-curated genomes belonging to Suli. Then, we focused on the remaining genomes. A set of 192 genomes with SS>4 were labeled as Sulfur unconsidered or related microorganisms (Sur). Finally, the rest of genomes in Gen were classified as NS (Non-Sulfur = Gen − (Suli + Sur)), including 1,754 genomes. The boxplots in Figure 5A summarize the scores obtained in these three subsets.

**Figure 5.**
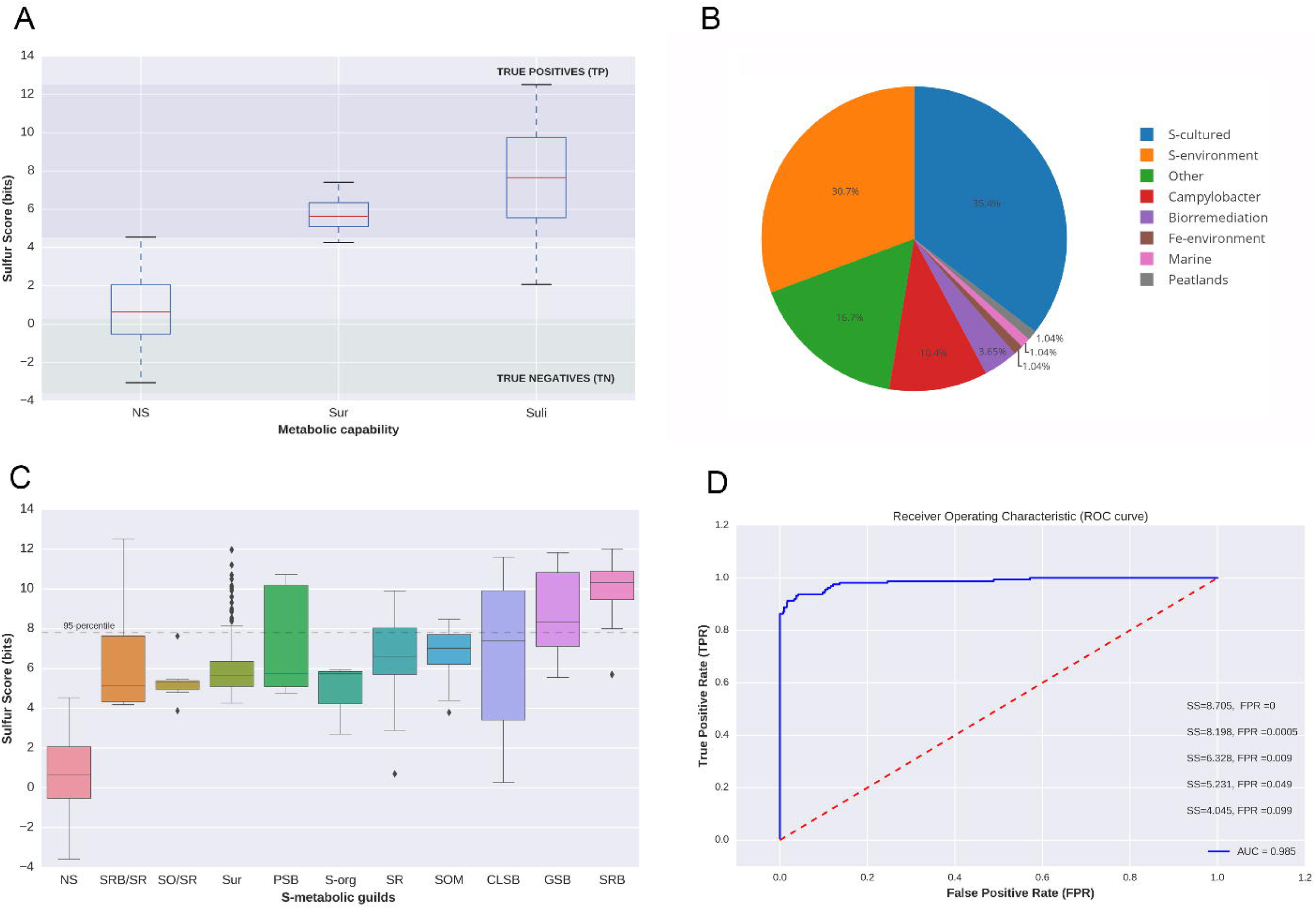
Distribution of Sulfur Score (SS) in 2,107 non-redundant genomes (Gen). A) Subsets of genomes annotated in Suli (n=161); ii) Sur, genomes not listed in Suli with SS > 4 and candidates to be S-related microorganisms (n=192); iii) rest of the genomes in Gen (NS, n=l,754). According to the curated species, True Positives can be defined as genomes with SS > max (SS_N5_) distribution, whereas True Negatives are those with SS < min(SS_Suli_). B) Assignment of the 192 genomes in Sur to ecological categories based on literature reports. C) Distribution of SS for different S-metabolic guilds, and the genomes in Sur. D) ROC curve with Area Under the Curve (AUC) indicated together with thresholds for some False Positive Rates (FPR).

To double-check whether the Sur genomes — selected due to their *SS* — might be involved in the S-cycle, we manually annotated all of them focusing on relevant genomic, biochemical, physiological and environmental information that we might have missed since Suli was first curated (Table S7). Out of 192 genomes, 68 are reported to metabolize S-compounds under culture conditions in the literature. For instance, *Sideroxydans lithotrophicus ES*-1, a microaerophilic Fe-oxidizing bacterium, has been observed to also grow in thiosulfate as an energy source [58]. Another 59 Sur organisms have been isolated from Sulfur-rich environments, such as hot springs or solfataric muds. Remarkably, some of this species include hard-to culture genomes reconstructed from metagenomic sequences such as *Candidatus Desulforudis audaxviator MP104C* isolated from basalt-hosted fluids of the deep subseafloor [6]; an unnamed endosymbiont of a scaly snail from a black smoker chimney [59] and archaeon *Geoglobus ahangari,* sampled from a 2,000m depth hydrothermal vent [60]. Furthermore, we also confirmed within Sur the implication of S-cycle of 20 species of the genus *Campylobacter.* These results are consistent with the ecological role of the involved taxa, that along with SRB and methanogens inhabiting host-gastrointestinal and low oxygen environments, where several inorganic (e.g., sulfates, sulfites) or organic (e.g., dietary amino acids and host mucins) are highly metabolized by these metabolic guilds [61]. The implication of *Campylobacter* species in the S-cycle is also supported by the fact that some of them have been isolated from deep sea hydrothermal vents [62]. The remaining species in Sur were classified in different categories, including bioremediation (7), Fe-environment (2), marine (2), peat lands (2) and other environments (32, see Figure 5B).

When the *SS* values of genomes in Sur are compared to the S-metabolic guilds represented in Suli (e.g PSB, SRB, GSB), it can be seen that they are indeed similar and clearly separated from the rest of NS genomes (Figure 5C). This strongly suggests that high scoring genomes are indeed ecologically and metabolically implicated in the S-cycle.

Finally, in order to quantify the capacity of the *SS* to accurately classify S-related microorganisms, we computed a Receiver Operator Characteristic (ROC) curve (for a detailed description of ROC curves see [63]). We thus defined genomes annotated in Suli as positive instances, and the rest as negative ones. The results are shown in Figure 5D, with an estimated Area Under the Curve (AUC) of 0.985, and the corresponding cut-off values of *SS* for several False Positive Rates (FPR). According to this test, a *SS value of* 8.705 is required to rule out all false positives in Gen, while SS=5.231 is sufficient to achieve a FPR < 0.05.

Overall, these results indicate that MEBS is a powerful and broadly applicable approach to predict, and classify microorganisms closely involved in the sulfur cycle even in hard-to culture microbial lineages.

### Sulfur Score and its predictive capacity to detect S-related environments in a large metagenomic collection

The SS was also computed for each metagenome in Met, using their corresponding MSL to choose the appropriate entropies previously calculated in dataset GenF (Table S8). In order to test whether *SS* values can be used to identify S-related environments, we performed the following analyses. First, we use the geographical metadata associated with each metagenome to map the global distribution of *SS*. In Figure 6A, *SS* values are colored from yellow to red. The most informative S-environments (displaying *SS* values equal or greater than the 95^th^ percentile of each MSL category) are shown in blue.

**Figure 6.**
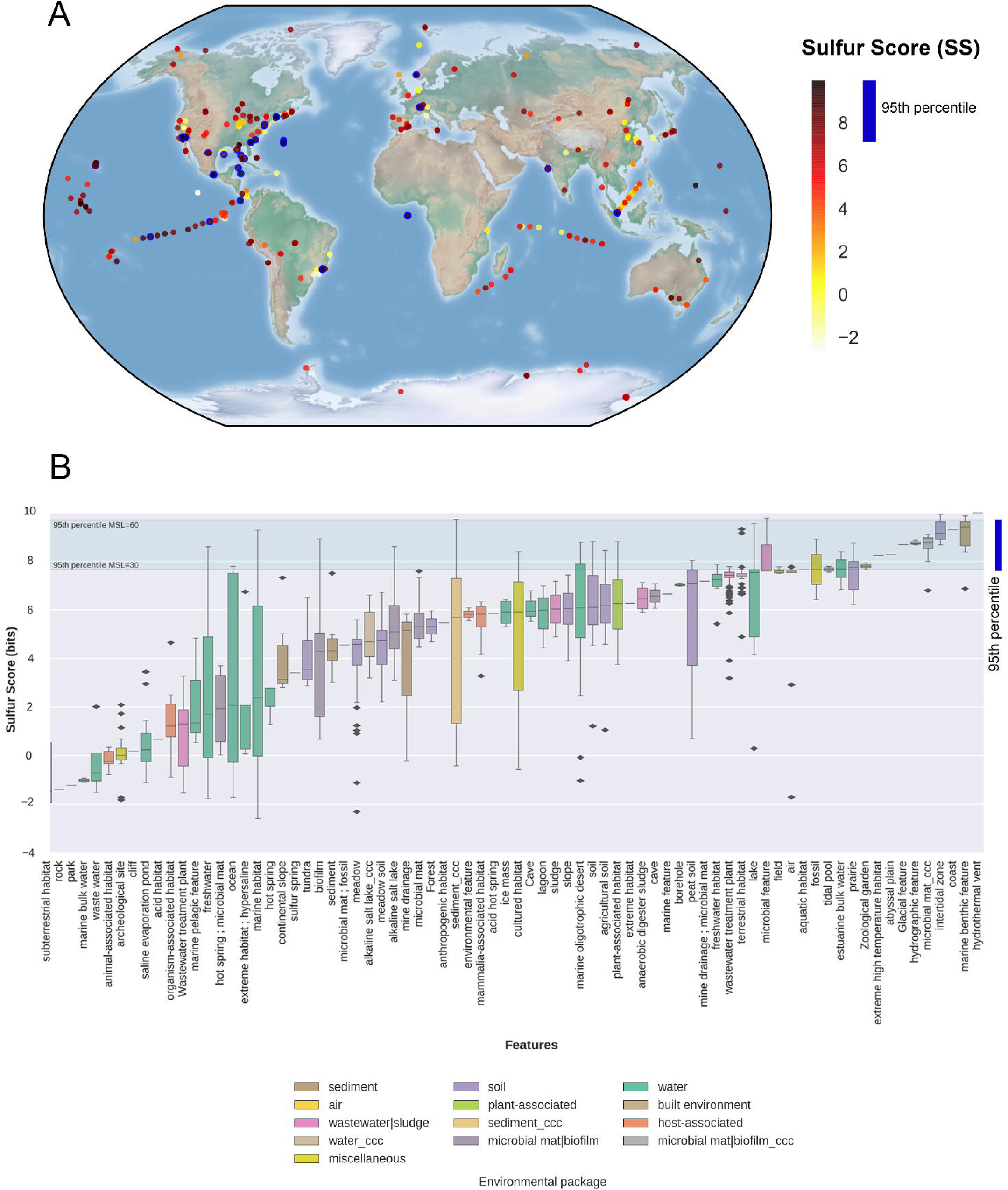
Distribution of Sulfur Score (SS) in the metagenomic dataset Met. A) Geo-localized metagenomes sampled around the globe are colored according to their *SS* values. The following cut-off values correspond to the 95th percentiles of seven Mean Size Length classes (30, 60, 100, 150, 200, 250 and 300 aa): 7.66, 9.70, 8.81, 8.51, 8.18, 8.98 and 7.61, respectively. Circles with thick blue border indicate metagenomes with SS ≥ the 95th percentile. B) Distribution of SS values observed in 935 metagenomes classified in terms of features (X-axis) and colored according to their particular habitats Features are sorted according to their median SS values, ccc: metagenomes from Cuatro Cienegas, Coahuila, Mexico. Green lines indicate the lowest and largest 95th percentiles observed across MSL classes.

Then, we sorted the metagenomes according to their environmental features as proposed by the Genomic Standards Consortium [GSC] and implemented in MG-RAST. Each feature corresponds to one of 13 environmental packages (EP) that standardize metadata describing particular habitats that are applicable across all GSC checklists and beyond [64]. Therefore, each EP represents a broad and general classification containing particular features. For example, the “water” EP includes 330 metagenomes from our dataset, belonging to several features such as freshwater, lakes, estuarine, marine, hydrothermal vents, etc. Since each of these features has different ecological capabilities in terms of biogeochemical cycles, we can expect different behaviors among *SS* values, as shown in Figure 6B. In general, all the metagenomes derived from hydrothermal vents (2), marine benthic (6), intertidal (8) and our unpublished CCC microbial mats had *SS* values above the 95^th^ percentile, highlighting the importance of the S-cycle in these environments. In contrast, the metagenomes belonging to features such as sub-terrestrial habitat (7), saline evaporation pond (24) or organisms associated habitat (7) displayed consistently low or even negative *SS* values, indicating a negligible presence of S-metabolic pathways in those environments. The remaining features have intermediate median *SS* values and contain occasionally individual metagenomes with *SS* values above the 95^th^ percentile, such as freshwater, marine, ocean or biofilm environments.

To validate the list of 50 high-scoring metagenomes (above the 95^th^ percentiles), we doublechecked their annotations. According to the literature and associated metadata, all these environments are closely involved in mineralization, uptake, and recycling processes of S-compounds. For example, environmental sequences derived from costal Oligochaete worm *Olavius algarvensis,* hydrothermal vents and marine deep-sea surface sediments around the Deep-Water Horizon spill in the Gulf of Mexico. The complete list of annotated metagenomes, along with their ecological capabilities, is found in Table S9.

### Evaluating the robustness of the Sulfur Score

To test the reproducibility and robustness of MEBS final score (*SS*), we conducted two further analyses. In the first one we compared *SS* estimates derived from Met dataset, computed with Pfam entropies obtained in the first MEBS benchmark performed three years ago (2014) with the current data described in this article (2017). Despite the changes of both databases (Pfam database version and the Suli list), we found a strong correlation (r^2^=0.912) between the *SS* outcomes (Figure S6A). A kernel density analysis of the latter comparison suggests a different behavior of low and high *SS* scores, with the latter being more reproducible (see Figure S6B).

In the second analysis, we quantitatively tested to what extent the entropy estimates of the 112 Pfam domains directly affect the outcome of the *SS* in Gen and Met. We randomly subsampled ≈50% of those domains to compute the *SS* a thousand times for each genome and metagenome in Gen and Met, respectively. The results, summarized in Table S10, confirm that *SS* values computed with random subsets of Pfam domains are generally lower than *SS* derived from the full list (n=112) of Sucy-Pfam domains. To further inspect the distribution of *SS* values produced with random subsets of domains (random *SS*), we focused on the particular case of the metagenomes belonging to the category MSL=60. As expected, the distribution of random *SS* oscillates between negative and positive values. Interestingly, metagenomes exhibiting only positive random *SS* are ranked above the 95^th^ percentile according to their real *SS* values (See Figure S7A). The latter indicates that even a random subset of Pfam domains are used to compute the score, is more likely to high-rank metagenomes containing the sulfur metabolic machinery (large number of high-entropy Pfam domains), than those lacking the sulfur metabolism or displaying a large number of non-informative Pfam domains. Furthermore, by comparing the median of random *SS* with the real scores, we observe a clear separation between those distributions (see Figure S7B and Table S10).

### Completeness of S-metabolic pathways

As we described above, the MEBS pipeline models a metabolic network as an array of S-related protein domains (Sucy-Pfam), to ultimately use their entropies to produce the final score (*SS*). For a closer look, we also dissected the total contribution of independent domains at the network level, in order to assess whether SS depends on the partial or complete detection of S-pathways. Consequently, we evaluated the pathway completene*SS* in both genomic (Gen) and metagenomic (Met) datasets (see Tables S11 and S12, respectively). Since the number of Pfam domains per pathway goes from one to 29 (see Table 1 and Table S2), we suspect that pathways represented by a single domain might not reflect their complete metabolic function. For example, the pathways involved in the methanogenesis of compounds such as dimethylsulfide (DMS, P24), methylthiolpropanoate (MTPA, P25), and methanethiol (MeSH, P26) are represented by the same protein (MtsA, PF01208) in our Sucy database, as well as in Metacyc [12]. Therefore, we expect that pathways P24-26 will have identical presence-absence patterns in Gen and Met.

The boxplots in Figure 7A and 7B summarize the distribution of completeness for each S-metabolic pathway including the synthetic pathway (P29) composed by 12 candidate markers as described in Table 2. As expected, the observed completeness per pathway was higher in Met than in Gen, since microbial communities harbor a wider repertory of metabolic functions than single genomes. In the case of genomes, we noted that a few pathways were complete in most genomes, being the majority involved in the usage of organic sulfur compounds such as alkanesulfonates (P9), sulfoacetate (P14) and biosynthesis of sulfolipids (SQDG) and the single domain pathways P24-26. Remarkably, we also detected a few organisms displaying the highest levels of metabolic completeness in some S-energy based pathways. For example, we found that *Desulfosporosinus acidiphilus SJ4* (SS=8,91) was the only genome harboring the complete repertory of Pfam domains described in Sucy for the sulfite oxidation (P1), strongly suggesting that it may oxidize sulfite. However, this activity remains to be tested in culture [65]. In the case of thiosulfate oxidation (P3), we detected three genomes displaying the highest levels of completeness, in agreement with their ecological features: *Hydrogenobaculum sp. Y04AAS1* (SS=9,319) [66] and the CLSB: *Acidithiobacillus caldus ATCC51756* (SS=6,525) [67] and *Acidithiobacillus ferrivorans* (SS=7,436) [68]. For the sulfate reduction dissimilative pathway (P5), out of 55 genomes displaying the higher completeness levels, 67% are actually SRB, 12% are Sur genomes, and the rest are sulfur oxidation microorganisms. Furthermore, the PSB *Thioflavicoccus mobilis 8321* (SS= 9,756), isolated from a microbial mat [69], was the genome displaying the most complete sulfide oxidation pathway (P11). Elemental sulfur disproportionation (P21) is represented by a single non-informative domain (PF07682, /-/’=0.172) that remarkably is found in 14 sulfur respiring or related genomes such as *Sulfolobus tokodaii str. 7* (SS= 5,341) and *Acidianus hospitalis W1* (SS= 3,88). Finally, we identified six genomes encoding all 12 proposed markers. Among them, three were GSB *(Pelodictyon phaeoclathratiforme BU-1,* SS=11,836, *Chlorobium chlorochromatii CaD3,* SS=11,625 and *Chlorobium tepidum TLS,* SS= 11,354), one CLSB (Thiobacillus denitrificans ATCC 25259 SS=11,61), another one PSB (*Thiocystis violascens DSM 198,* SS=10,633) and finally one Sur *(Sedimenticola thiotaurini* SS=10,109). For a complete description, see Table S13.

**Figure 7.**
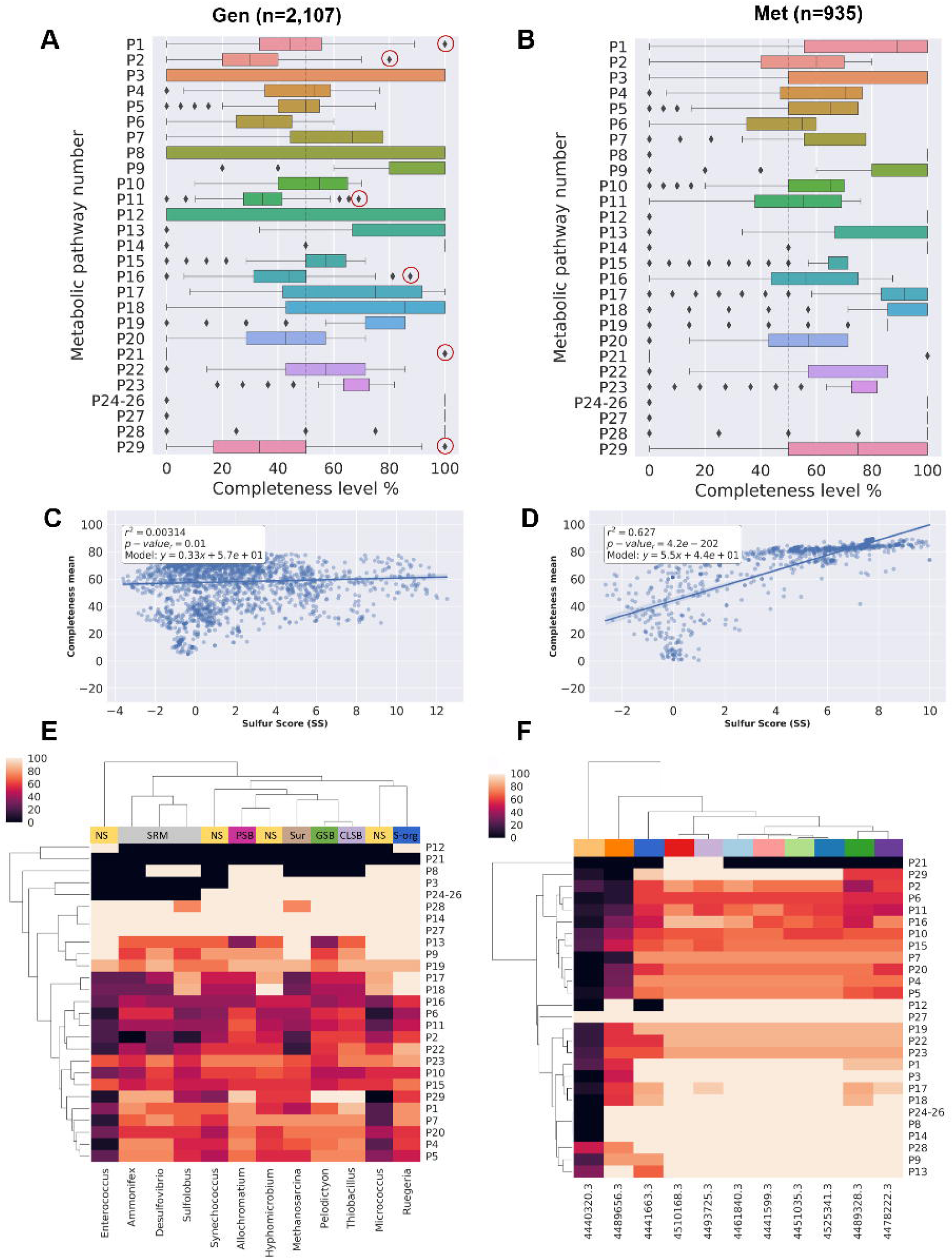
Metabolic completeness of the metabolic pathways described in Table 1. A) Boxplot distribution of the pathway completeness in genomic and B) metagenomic datasets. C) Linear regression models of the Sulfur Score (SS) and the mean completeness in Gen and D) Met dataset. E) Heatmap showing the metabolic completeness of the following genomes: *Desulfovibrio vulgaris DP4* (55=11,442), *Ammonifex degensii KC4* (SS=12.508); *Pelodictyon phaeoclathratiforme BU-1* (55=11,836); *Thiobacillus denitrificans ATCC 25259* (SS=11,61); PSB: *Allochromatium vinosum DSM 180* (SS=10. 737); Sur Methanosarcina barkeri MS(SS= 5,93); *Sulfolobus acidocaldarius DSM 639* (SS=5,457); *Synechococcus sp. JA-2-3Ba 2-13* (SS=3,704); *Hyphomicrobium denitrificans 1NES1 {SS-* 3,236); *Ruegeria pomeroyi DSS-3 (SS-2,707)·, Enterococcus durans* (SS=-0,194); *Micrococcus luteus NCTC_2665* (SS=-3,588). F) Heatmap showing the metabolic completeness of the metagenomes with the following MG-RAST ids and corresponding scores: 4489656.3 (SS=-2,649); 4440320.3(SS=0,1); 4441663.3(SS=9,986); 4510168.3 (SS=7,781); 4493725.3 (SS=9,547); 4461840.3 (SS=8,813); 4441599.3(SS=9,274); 4451035.3(SS=9,918); 4525341.3(SS=9,287); 4489328.3(SS=4,958); 4478222.3(SS=4,88). The color codes at the top of the heatmap correspond to different environments. For a more detailed description of each metagenome see Table S8.

A global view of metabolic completeness was obtained by bulking the data from all pathways. Linear regression models between mean completeness and *SS* were computed confirm the, yielding r^2^ values of 0.003 and 0.627 for Gen and Met, respectively (See Figures 7C and 7D). Moreover, we also assessed the relationship between the mean completeness of the synthetic pathway of candidate markers (P29) and the SS. As expected, significant correlations were obtained in both datasets (r^2^= 0.645 and r^2^=0.881 for Gen and Met, respectively; see Figure S8).

To get a more detailed insight of the completeness, we selected a few genomes and metagenomes displaying high and low *SS* values. Specifically, from the Gen dataset we selected one representative from the main S-guilds, one Sur genome and two genomes with low SS values (NS). As observed in Figure 7, the low-scoring genomes *Enterococcus durans* (SS=-0,194), *Micrococcus luteus NCTC_2665* (SS=-3,588), *and Ruegeria pomeroyi DSS-3* (SS=2,707) display unrelated patterns of sulfur metabolic completeness, compared with the rest of genomes and therefore are separated. In contrast, high-scoring S-respiring microorganisms *Desulfovibrio vulgaris DP4* (SS= 11,442), *Sulfolobus acidocaldarius DSM 639* (SS=5,457) and *Ammonifex degensii KC4* (SS=12.508) are clustered together. We also observed that mat-isolated cyanobacteria *Synechococcus sp. JA-2-3Ba 2-13,* classified as NS with SS=3,704, was clustered together with other high-scoring genomes, in agreement with the lack of correlation reported above.

In the case of metagenomes (see Figure 7E), we observed a clear correlation between SS and completeness. For example, metagenomes 4440320.3 and 4489656.3, with the lowest scores (SS=0.1 and SS=-2.649, respectively), also exhibit the largest number of incomplete pathways. Similarly, high-scoring metagenomes derived from black smoker or marine sediment are grouped together in terms of completeness.

## Conclusions

Our study represents the first exploration of the Sulfur biogeochemical cycle in a large collection of genomes and metagenomes. The manually curated effort resulted in an inventory of the compounds, genes, proteins, molecular pathways, and microorganisms involved. This complex universe of articulated data was reduced into a list of microorganisms and Pfam domains encoded in the proteins that take part in that network. These domains were first ranked in terms of relative entropy, and then summed to produce a single S-score representing the relevance of a given genomic or metagenomic sample in terms of sulfur metabolic machinery. We took advantage of the mathematical framework of information theory, which has been widely used in computational biology.

The performance of the Multigenomic Entropy Based Score pipeline (MEBS) (designed for the above mentioned tasks) was benchmarked on large genomic and metagenomic sets. Our results support the broad applicability of this algorithm in order to classify annotated genomes as well as newly sequenced environmental samples without prior culture. We also assessed to what extent the final score depended on the partial or complete detection of pathways and observed a higher completeness per pathway in metagenomic sequences than in individual genomes.

We demonstrated that a measurable score can be applied to evaluate any given metabolic machinery or biogeochemical cycle in large (meta)genomic scale, holding the potential to dramatically change the current view of inferring metabolic capabilities in the present ‘omic’-era.

## Availability and requirements

Project name: MEBS

Project home page: https://github.com/eead-csic-compbio/metagenome_Pfam_score

Operating system(S): Linux

Programming language: Python 3, Perl5, Bash,

Other requirements: HMMER

License: GNU General Public License (GPL)

## Availability of supporting data

The datasets supporting the results of this article are available in the GigaDB repository [REF#]

## Abbreviations

MEBS: Multigenomic Entropy Based Score
S: Sulfur
S-cycle: Sulfur cycle
SS: Sulfur Score
Suli: Sulfur list
Sucy: Sulfur cycle database
Rlist: Random list of taxonomic representatives
MSL: Mean Size Length
H’: Relative Entropy
Sur: Sulfur unconsidered
NS: Non sulfur related genomes
Gen: Genomic dataset
Met: Metagenomic dataset
GenF: Genomic Fragmented dataset
CLSB: Color-less Sulfur Bacteria
SOM: Sulfur Oxidizing Microorganims
GSB: Green Sulfur Bacteria
PSB: Purple Sulfur Bacteria
SRB: Sulfate Reducing Bacteria
ESR: Elemental-Sulfur Reducing microorganisms
CCC: Cuatro Cienegas, Coahuila
HMM: Hidden Markov Models (HMMs)
ROC: Receiver-operating characteristic
AUC: Area Under the Curve
TPR: True Positive Rates
FPR: False Positive Rates
GSC: Genomic Standards Consortium
EP: environmental packages

## Acknowledgments

The authors gratefully acknowledge Emilio Morella for their valuable support, feedback and comments throughout the development of this project. We really appreciate the reviewers Daan Speth and Thulani Makhalanyane whose valuable comments and suggestions critically improve the manuscript and sofware. Our special thanks to acknowledge Carlos P Cantalapiedra, Seth Barribeau, Will Levitt and anonymous reviewers for their comments on earlier versions of the manuscript who immensely improved the algorithm and the final version of the article. The authors also thank the Laboratory of Computational and Structural Biology (EEAD-CSIC) and Laboratorio de Evolución Molecular y Experimental, Instituto de Ecología, UNAM for providing the computational resources described in the article. The paper was written during a sabbatical leave of LEE and VSS in the University of Minnesota in Peter Tiffin and Michael Travisano laboratories.

## Funding

VDA is a doctoral student from Programa de Doctorado en Ciencias Biomédicas, Universidad Nacional Autónoma de México (UNAM) and received fellowship 356832 from CONACYT. This research was also supported by funding from WWF-Alianza Carlos Slim, Sep-Ciencia Básica Conacyt grant 238245 to both VS and LEE and Spanish MINECO grant CSIC13-4E-2490. BCM was funded by Fundación ARAID. The sabbatical leave of LEE and VSS at the University of Minnesota were supported by scholarships from PASPA, DGAPA, UNAM.

## Competing interest

The authors declare that they have no competing interest.

## Author contribution

VDA, BCM and IZP wrote the paper. BCM, VDA and ACPH developed and wrote the software and performed all the bioinformatics analyses. VDA produced all the figures and wrote the documentation of the software. VDA and IZP conceived the manual curation of the Sulfur cycle inventory and the microbiological, biogeochemical, and ecological interpretation. LE and VS provided the intellectual framework, expertise and resources to develop and supervise the project. All the authors read and approved the final manuscript.

## Endnotes

We are currently finishing the analyses to demonstrate the applicability of this approach to other biogeochemical cycles (C, N, O, Fe, P). Thereby, we hope that the pipeline MEBS will facilitate analysis of biogeochemical cycles or complex metabolic networks carried out by specific prokaryotic guilds, such as bioremediation processes (i.e., degradation of hydrocarbons, toxic aromatic compounds, heavy metals etc.). We look forward to collaborate and help other researchers by integrating comprehensive databases that might be helpful to the scientific community. Furthermore, we are currently working to improve the algorithm by using only a list of sequenced genomes involved in the metabolism of interest, in order to reduce the manual curation effort. We are also considering taking *k-mers* instead of peptide Hidden Markov Models to increase the speed of the pipeline. We anticipate that our platform will stimulate interest and involvement among the scientific community to explore uncultured genomes derived from large metagenomic sequences.

## Additional files-Supplementary Information

The supplementary pdf file contains the following information:

**Supplementary figure S1.** Histogram distribution of the Mean Size Length of metagenomes in Met and the input sulfur proteins.

**Supplementary figure S2.** Visualization of the Pfam domains mapped onto KEGG metabolic pathways

**Supplementary figure S3.** Comparison of clustering methods of the 112 Pfam entropies using script *plot_cluster_comparison.py*

**Supplementary figure S4.** Clustering comparison between Birch and Ward clustering methods to stand out the Pfam entropies with high *H’* and low std using the script

**Supplementary figure S5.**. Distribution of entropy values of 112 Pfam domains inferred from random-sampled and Suli genomes.

**Supplementary figure S6.** Comparison of Sulfur Scores (SS) with data obtained three years ago (2014), with the current data described in the article.

**Supplementary Table S4.** Informative Pfam’s with high *H’* and high std (not used as molecular marker genes) in metagenomic fragmented data.

**Supplementary Table S5.** Percentile distribution of the 112 Pfam entropies in the random test

**Supplementary Table S10.** Statistics of 55 computed on genomic (Gen, real sequences) and metagenomic (Met, with increasing Mean Size Length, from 30 to 300aa) datasets

In separated excel files the following Supplementary tables are also provided:

**Supplementary Table SI:** Table S1. Comprehensive list of the taxonomic representatives of sulfur cycle including Sulfur list or ‘Suli’ containing 161-curated genomes used as input for the pipeline

**Supplementary Table S2.** Sucy database containing the identifiers of the Sulfur proteins and their corresponding annotations derived from Interproscan and manual curation.

**Supplementary Table S3.** Sulfur Pfam domains (Pfam-Sucy), and their corresponding mapping into KEGG (KO number), and the manual assignation into sulfur metabolic pathways

**Supplementary Table S6.** Gen dataset containing their corresponding SS and taxonomy assignment.

**Supplementary Table S7.** Manual annotation of Sulfur unconsidered or related microorganisms (Sur) with SS>4 Supplementary Table S8

**Supplementary Table S8.** Met dataset with their corresponding SS values and metatada.

**Supplementary Table S9.** Manually annotated high scoring metagenomes along with their ecological capabilities in terms of sulfur cycle

**Supplementary Table S11.** Metabolic completeness in Gen dataset for each of the 28 metabolic pathways of the S-cycle described in Table 1. (Pathway 29 contains the proposed marker genes)

**Supplementary Table S12.** Metabolic completeness in Men dataset for each of the 28 metabolic pathways of the S-cycle described in Table 1. (Pathway 29 contains the proposed marker genes)

**Supplementary Table S13.** Frequency and description of the most complete genomes in terms of S-cycle metabolic pathways

